# On the structure of neuronal population activity under fluctuations in attentional state

**DOI:** 10.1101/018226

**Authors:** Alexander S. Ecker, George H. Denfield, Matthias Bethge, Andreas S. Tolias

**Affiliations:** Centre for Integrative Neuroscience and Institute for Theoretical Physics, University of Tübingen, Germany; Max Planck Institute for Biological Cybernetics, Tübingen, Germany; Bernstein Centre for Computational Neuroscience, Tübingen, Germany; Department of Neuroscience, Baylor College of Medicine, Houston, TX, USA; Department of Computational and Applied Mathematics, Rice University, Houston, TX, USA

## Abstract

Attention is commonly thought to improve behavioral performance by increasing response gain and suppressing shared variability in neuronal populations. However, both the focus and the strength of attention are likely to vary from one experimental trial to the next, thereby inducing response variability unknown to the experimenter. Here we study analytically how fluctuations in attentional state affect the structure of population responses in a simple model of spatial and feature attention. In our model, attention acts on the neural response exclusively by modulating each neuron’s gain. Neurons are conditionally independent given the stimulus and the attentional gain, and correlated activity arises only from trial-to-trial fluctuations of the attentional state, which are unknown to the experimenter. We find that this simple model can readily explain many aspects of neural response modulation under attention, such as increased response gain, reduced individual and shared variability, increased correlations with firing rates, limited range correlations, and differential correlations. We therefore suggest that attention may act primarily by increasing response gain of individual neurons without affecting their correlation structure. The experimentally observed reduction in correlations may instead result from reduced variability of the attentional gain when a stimulus is attended. Moreover, we show that attentional gain fluctuations – even if unknown to a downstream readout – do not impair the readout accuracy despite inducing limited-range correlations.

## Introduction

Attention was traditionally thought of as acting by increasing response gain of a relevant population of neurons (Maunsell and Treue 2006; Reynolds and Chelazzi 2004). More recent studies found that attention also reduces pairwise correlations between neurons (Cohen and Maunsell 2009; Herrero et al. 2013; Mitchell et al. 2009). Based on a simple pooling model (Zohary et al. 1994) these authors argued that the effects of increased gain are dwarfed by the effects of reduced correlations and, therefore, attention is more appropriately viewed as shaping the noise distribution.

However, in an experiment the subject’s state of attention can be controlled only indirectly and is bound to vary from one trial to the next. As a consequence, measuring neuronal variability or correlations under attention has a fundamental caveat: it is unclear to what extent the observed neuronal covariability reflects interesting aspects of information processing in the neuronal population or simply trial-to-trial fluctuations in the subject’s state of attention, which is unknown to the experimenter. Despite ample evidence that attention fluctuates from trial to trial (Cohen and Maunsell 2010; Cohen and Maunsell 2011), the effects of such fluctuations on neuronal population activity have so far not been investigated.

Here we analyze a simple neural population model, where neurons with overlapping receptive fields encode the direction of motion of a stimulus (Fig. 1A). We assume that neurons produce spikes independently according to a Poisson process with rate *λ_i_* and treat attention as a process that modulates the neurons’ gain (Fig. 1B). The firing rates are given by

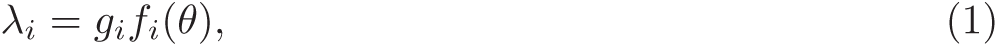

where *g*_*i*_ is the attentional gain (a combination of spatial and feature attention) and *f*_*i*_(*θ*) is the direction tuning curve of neuron *i*. We assume that there is always a stimulus in the neurons’ receptive field, but this stimulus is not necessarily attended. Crucially, in our model the subject’s attentional state is not constant across trials, even within the same attentional condition. Thus, *g*_*i*_ is a random variable that varies from trial to trial (Fig. 1C), and its precise value is unknown to the experimenter. As a consequence, the correlations in *g*_*i*_ across neurons will induce correlations between the observed neural responses.

**Figure 1:**
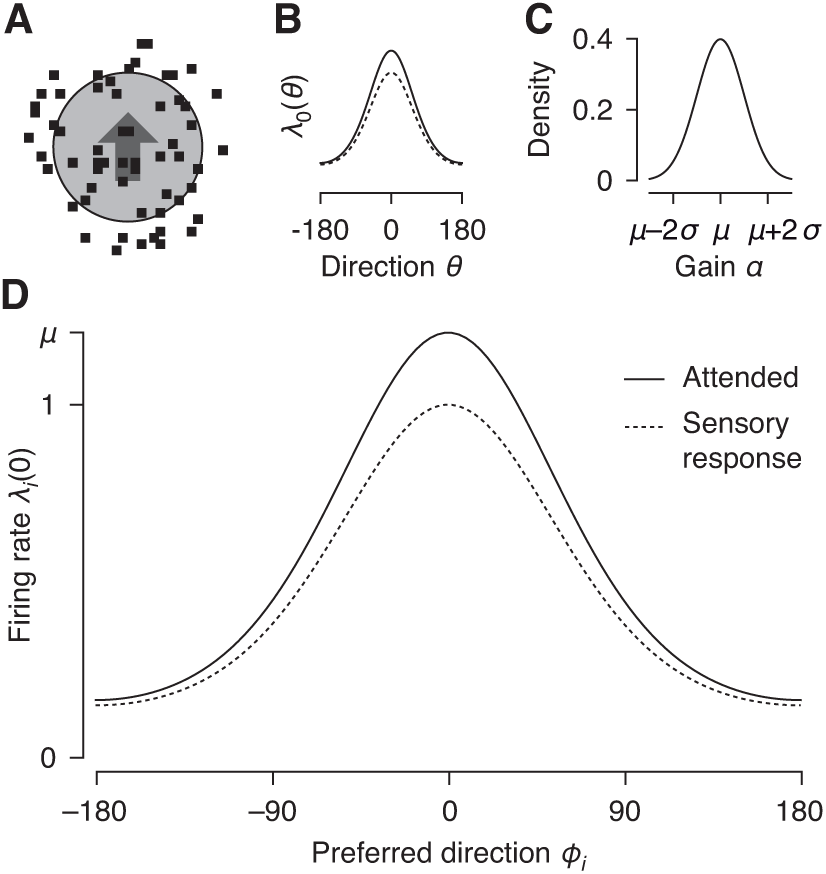
Model of spatial attention. **A.** Example stimulus. Neurons’ receptive fields are assumed to be at the same location (circle). **B.** Tuning curve under sensory stimulation (dashed) and with spatial attention directed to the stimulus in the receptive field (solid). **C.** Distribution of attentional gain (*α*). **D.** Population response of a homogeneous population of neurons under sensory stimulation (dashed) and with attention directed to the stimulus in the receptive fields (solid).

In the following sections, we analyze this correlation structure in detail. We find that the correlations induced by attentional fluctuations resemble many experimentally observed aspects of correlated variability, such as correlations that increase with firing rates, limited range correlations, and differential correlations. In addition, we investigate the consequences of correlations induced by fluctuating attentional gain for reading out the direction of motion of the stimulus from the population response. We show that such correlations do not impair readout, even if the decoder does not have access to the attentional state.

Our results have been presented previously in abstract form (Ecker et al. 2012). Some of the ideas presented in this paper have recently been developed independently by another group (Rabinowitz et al. 2015).

## 2 Methods

This section contains a detailed description of the model and the derivations of the main results. In an effort to make the paper as accessible as possible, the results section is self-contained. Readers not interested in the detailed derivations can skip ahead directly to the results section on page 8.

### 2.1 Model setup

We model a population of direction-selective neurons with overlapping receptive fields and a diverse range of preferred directions *Φ_i_*. We use a simple model of spatial and feature attention, where a neuron’s firing rate *λ*_*i*_ is the product of an attentional gain *g*_*i*_(*Ψ*) and a tuning function *f*_*i*_(*θ*):

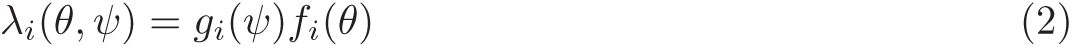

Here, *Ψ* is the attended direction of motion and *θ* the direction of the stimulus that is shown. Neurons are assumed to be conditionally independent given the firing rate *λ*_*i*_ (i.e. no noise correlations). The attentional gain depends on whether attention is directed to the location of the neurons’ receptive fields and on the attended direction of motion. For spatial attention, we use *g*_*i*_ = *α*, which is the same for all neurons, since they all have overlapping receptive fields. For feature attention we use *g*_*i*_(*Ψ*) = 1 + *βh*(*Ψ* - *Φ*_*i*_), where *β* the *feature attention gain*, and *h*(.) the *gain profile*. We follow the feature similarity gain model (Treue and Martinez-Trujillo 1999), where a neuron’s gain is enhanced if the attended feature matches the neuron’s preference and suppressed otherwise. A common choice for *h* is a cosine: *h*(*Ψ* - *Φ*_*i*_) = cos(*Ψ* - *Φ*_*i*_).

Note that from the perspective of the model there is no fundamental difference between spatial and feature attention. If we treat space as a variable that is being encoded by the population, any derivations for feature attention also apply to spatial attention. However, because we consider only a local population with overlapping receptive fields, spatial attention is a special case: the gain profile within the population is constant and therefore spatial attention can be expressed in a simpler way using a single common gain *α*. Thus, whenever we refer to spatial attention, this applies to a situation where all neurons in the population that is being considered share the same preferred feature. Likewise, whenever we refer to feature attention, this applies to any situation where the neurons in the population span a large range of preferred features. We chose this (somewhat arbitrary) distinction, because it reflects the typical situation in an experiment, where neurons with similar retinotopic locations are recorded, which typically span a large range of preferred orientations or directions.

### 2.2 Effect of fluctuating gains on spike count statistics

Throughout this paper we assume that spatial and feature attention are independent processes and consider them in isolation. We further assume that the experimenter does not have access to the attentional state on individual trials, but can only control its average over many trials:

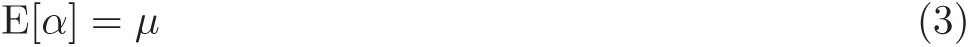

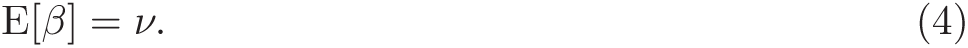

In addition the attentional state fluctuates from trial to trial with unknown variance

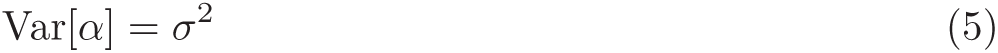

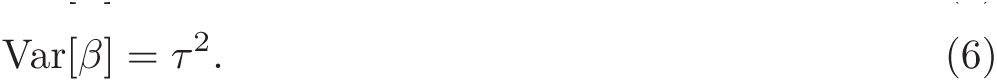

To compute means and (co-)variances of the observed spike counts we need only the means and variances of *α* and *β*. The expected spike counts (Eqs. 31, 37) follow from the linearity of the expectation. Variances and covariances can be computed by application of the Law of Total Variance (here for the case of spatial attention, feature attention follows the same logic):

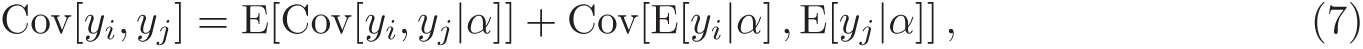

where the outer expectation (covariance) is taken over *α* and the inner covariance (expectation) over *y*_*i*_ and *y*_*j*_. Plugging the definitions of *λ*_*i*_ = E[*y*_*i*_|*α*] and using the assumption of conditionally independent Poisson spiking Cov[*y*_*i*_, *y*_*j*_|*α*] = *δ_ij_λ_i_*, we obtain the expressions for variances and covariances stated in the Results (Eqs. 32–34, 38–41).

### 2.3 Effect of fluctuations in attended feature on spike count statistics

Calculating the means and covariances under fluctuations in the attended direction *Ψ* follows the same approach as above. However, since the gain profile *h*_*i*_(*Ψ*) can be non-linear, we need a few additional assumptions. We assume that *Ψ* is distributed around some direction *Ψ*_0_ = E[*Ψ*] with variance *q*^2^ = Var[*Ψ*]. For reasonably small *q*^2^ we can approximate the gain profile by its first-order Taylor expansion

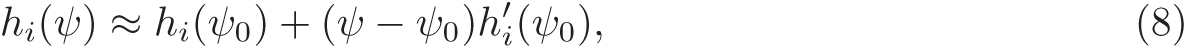

where 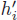 is the derivative with respect to *Ψ*. Using this approximation we can write E[*h*_*i*_(*Ψ*)] ≈ *h*_*i*_(*Ψ*_0_) and 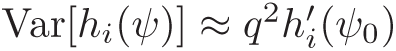, which leads (again after applying the Law of Total Variance) to the results in Eqs. 42–44.

### 2.4 Coding accuracy under fluctuations of spatial attention

Here we show that fluctuations in spatial attention have a negligible effect on the amount of information about the orientation of the stimulus. For simplicity we assume that neurons produce spikes conditionally independently given the stimulus orientation *θ* and the attentional gain *g*:

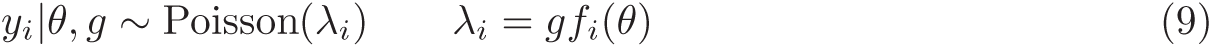

The attentional gain *g* is shared among all neurons and drawn from a Gamma distribution with shape *μ*^2^*/σ*^2^ and scale *σ*^2^*/μ*, which implies E[*g*] = *μ* and Var[*g*] = *σ*^2^. Assuming that the experimenter does not know the attentional gain, the distribution *P* (y|*θ*) obtained by marginalizing over *g* is a multivariate negative binomial distribution:

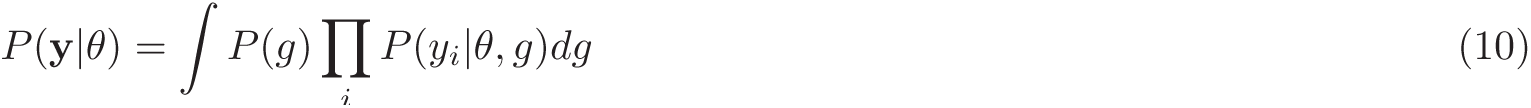

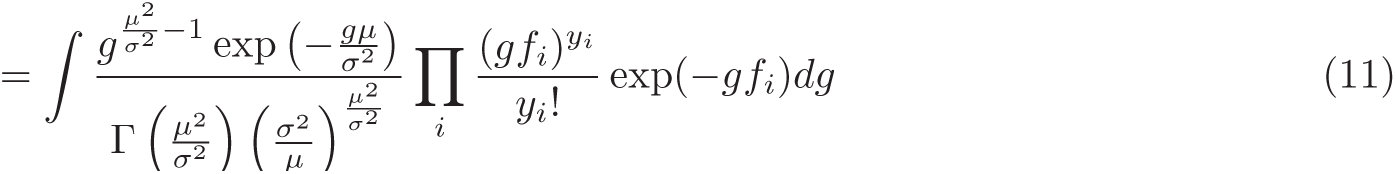

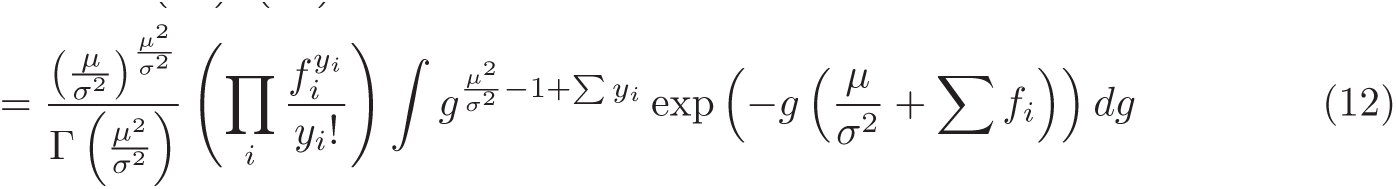

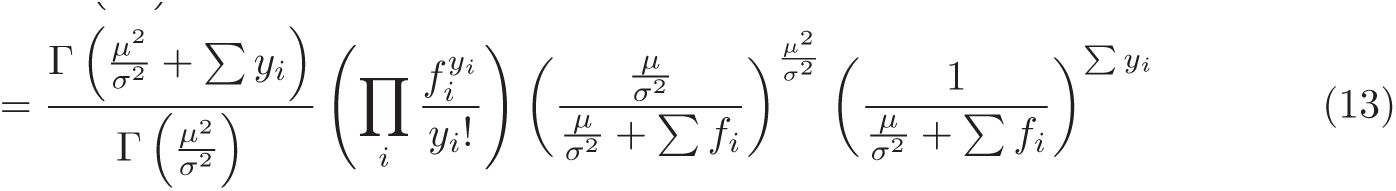

For the Fisher information 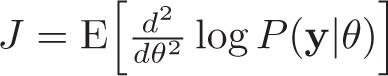 we need the derivatives of the log-likelihood:

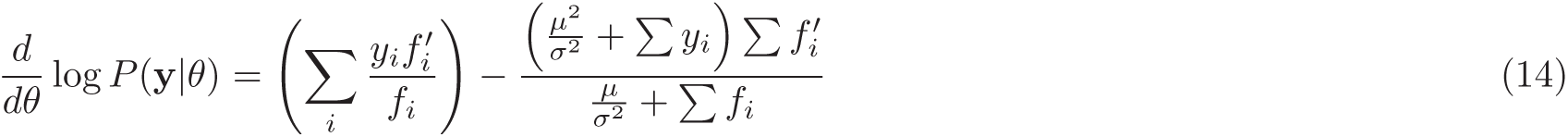

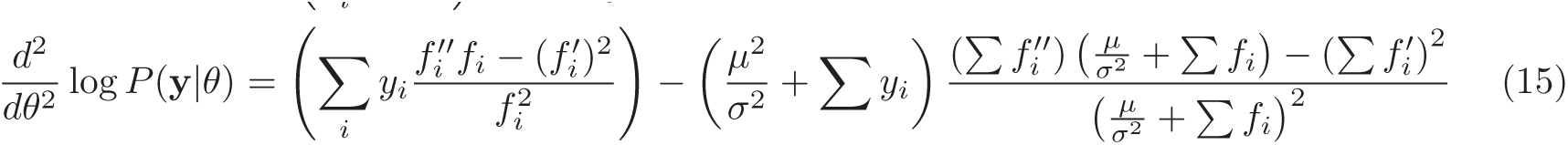

Plugging into the formula for Fisher information, re-ordering the summations over y and *i*, and using the facts ∑_y_*P* (y|*θ*) = 1 and ∑_y_ *P* (y|*θ*)*y*_*i*_ = E[*y*_*i*_] = *μf_i_*, we obtain

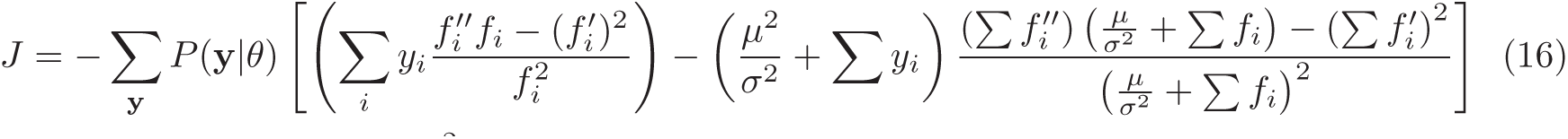

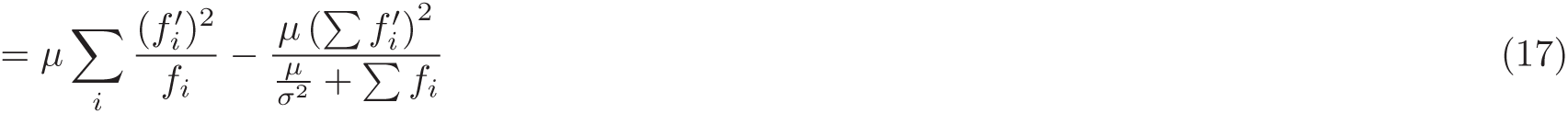

The first term in the above equation is the Fisher information of an independent population of neurons and therefore *O*(*N*), while the second term is *O*(1): for homogeneous population of neurons, where *f*_*i*_(*θ*) = *f* (*θ* - *Φ*_*i*_), it is zero; for heterogeneous populations it is *O*(1), as we show in the next paragraph. Thus, fluctuations in spatial attention do not impair the coding accuracy of the population with respect to orientation.

To show that the second term above is *O*(1) for heterogeneous populations, we assume that the neurons’ tuning curves are independent random variables (see Ecker et al. 2011; Shamir and Sompolinsky 2006). In this case the quantity of interest is the expected value with respect to different realizations of the heterogeneity:

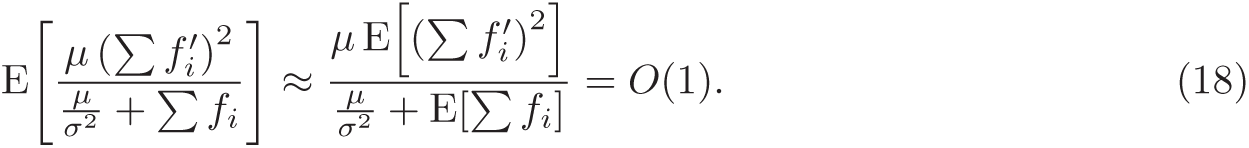

Here the approximation holds because for large *N* the width of the distribution of ∑ *f*_*i*_ becomes narrower relative to its mean and therefore the expected value of the second term converges to the ratio of the expected values of numerator and denominator. The equality holds because ∑ *f*_*i*_ = *O*(*N*) and

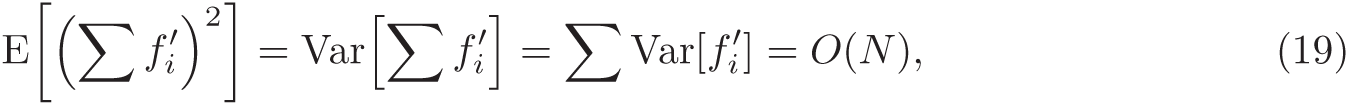

which holds because 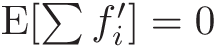.

### 2.5 Coding accuracy under fluctuations of feature attentional gain

Fluctuations in feature attention are more difficult to study analytically. Unfortunately, the Gamma-Poisson mixture model employed above does not generalize to the case where the gain is weighted differently for each neuron (i.e. the gain profile *h*_*i*_), or at least we are not aware of a model that has a closed-form expression for the marginal probability mass function when the gain is unknown. Therefore, we here approximate the population activity by a multivariate Gaussian distribution with matching mean and covariance matrix (Eq. 37–41) and focus on linear readout. Under this approximation, the (linear) Fisher Information is given by

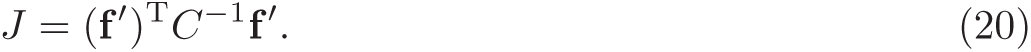

The inverse of the covariance matrix is obtained by applying a rank-one update:

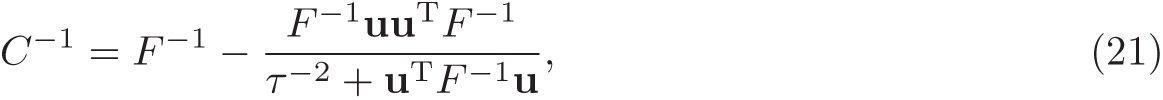

where *F*_*ii*_ = (1 + *νh_i_*(*Ψ*))*f*_*i*_(*θ*) and *u*_*i*_ = *h*_*i*_(*Ψ*)*f*_*i*_(*θ*) as above. Plugging in and simplifying we obtain

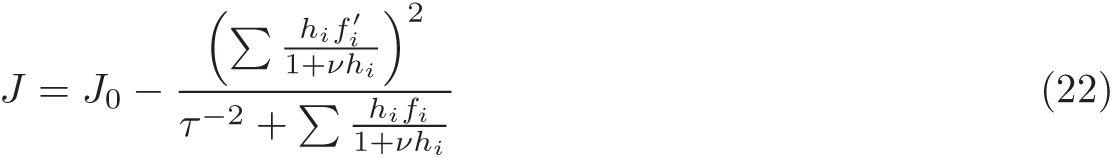

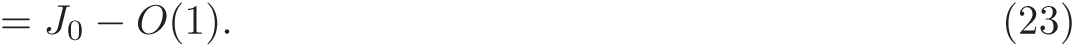

As above for spatial attention, the *O*(1) correction term is exactly zero for homogeneous populations and the derivation for heterogeneous populations follows the same line of argument as above.

### 2.6 Coding accuracy under fluctuations of attended feature

Fluctuations of the attended feature create differential correlations, i. e. response variability that is identical to variability induced by changes in the stimulus. Here we derive this result using a Generalized Linear Model formulation (see also Eqs. 52, 53 in Results):

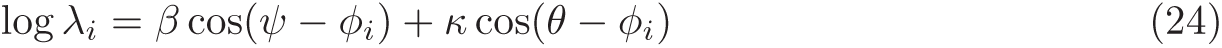

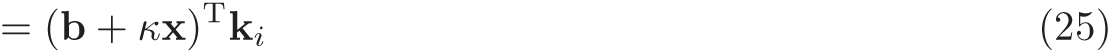

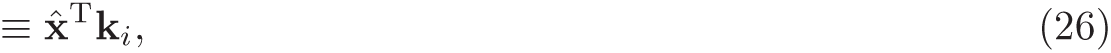

where b = *β*[cos *Ψ,* sin *Ψ*]^T^, x = [cos *θ,* sin *θ*]^T^, and *k*_*i*_ = [cos *Φ_i_,* sin *Φ*_*i*_]^T^. Since 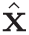 is independent of the neurons, it is obvious that attention has exactly the same effect as a change in the stimulus. Assuming E[*Ψ*] = *θ*, Var[*Ψ*] is small, and (without loss of generality) *θ* = 0, we have

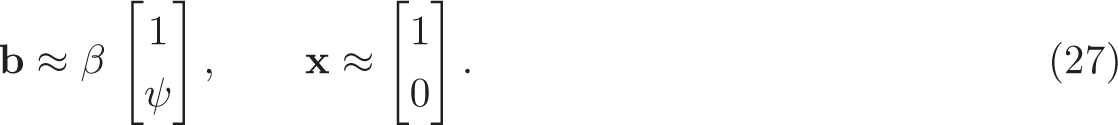

Moreover, we can write the attention-perturbed stimulus 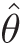 as

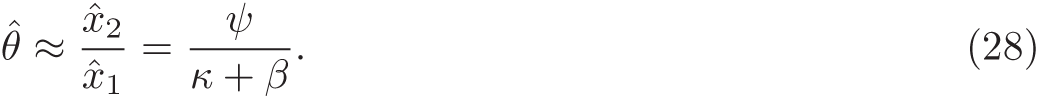

For large *N* the Poisson noise averages out and therefore the resulting Fisher information is simply the inverse of the variance of the (attention-perturbed) stimulus:

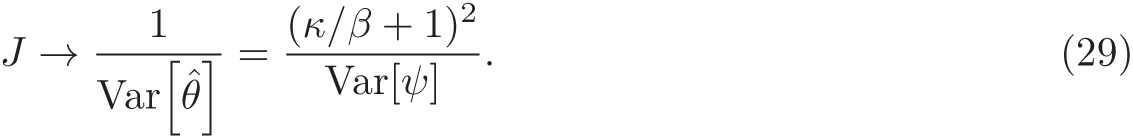

### 2.7 Code

Figures were generated using Matlab R2014b (The Mathworks Inc.). The code to reproduce the figures is publicly available at https://github.com/aecker/attentional-fluctuations.

## 3 Results

### 3.1 Fluctuations in spatial attention

Our goal is to characterize the effect of fluctuating attentional signals on the population response in sensory areas. We start by considering the simplest case of spatial attention and a common gain *α* for all neurons (Fig. 1):

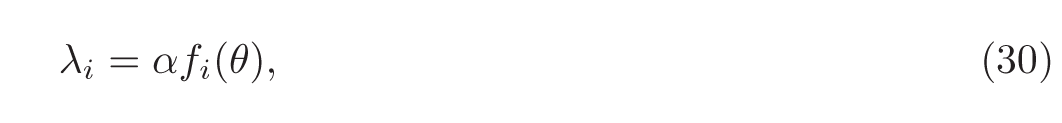

where *α >* 0 is the amount of spatial attention allocated to the stimulus in the neurons’ receptive field. We do not require any distributional assumptions on *α*, except for its mean E[*α*] = *μ* and variance Var[*α*] = *σ*^2^ (Fig. 1C). Under this model, the average spike count of a neuron is given by

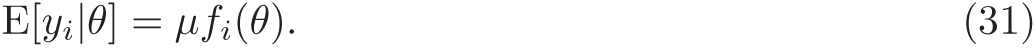

By convention we refer to the case of *μ* = 1 as the *sensory response*, which is the neural response to the stimulus in the absence of any attentional modulation. In experimental conditions where the stimulus is attended *μ_a_ >* 1 (Fig. 1D). When attention is directed towards a different stimulus *μ*_*u*_ ≤ 1 (depending on whether responses are suppressed relative to the sensory response under such conditions). Note that although we use homogeneous neural populations in the figures (all neurons have the same tuning curve up to a preferred direction *Φ*_*i*_, i.e. *f*_*i*_(*θ*) = *f* (*θ* - *Φ*_*i*_)), all results hold more generally for arbitrary tuning curves.

Because the attentional state fluctuates from trial to trial, the underlying firing rate also fluctuates. By applying the law of total variance we obtain the spike count variance (Fig. 2A):

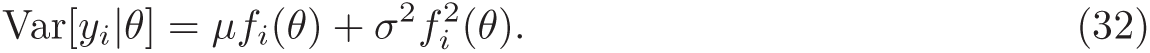

**Figure 2:**
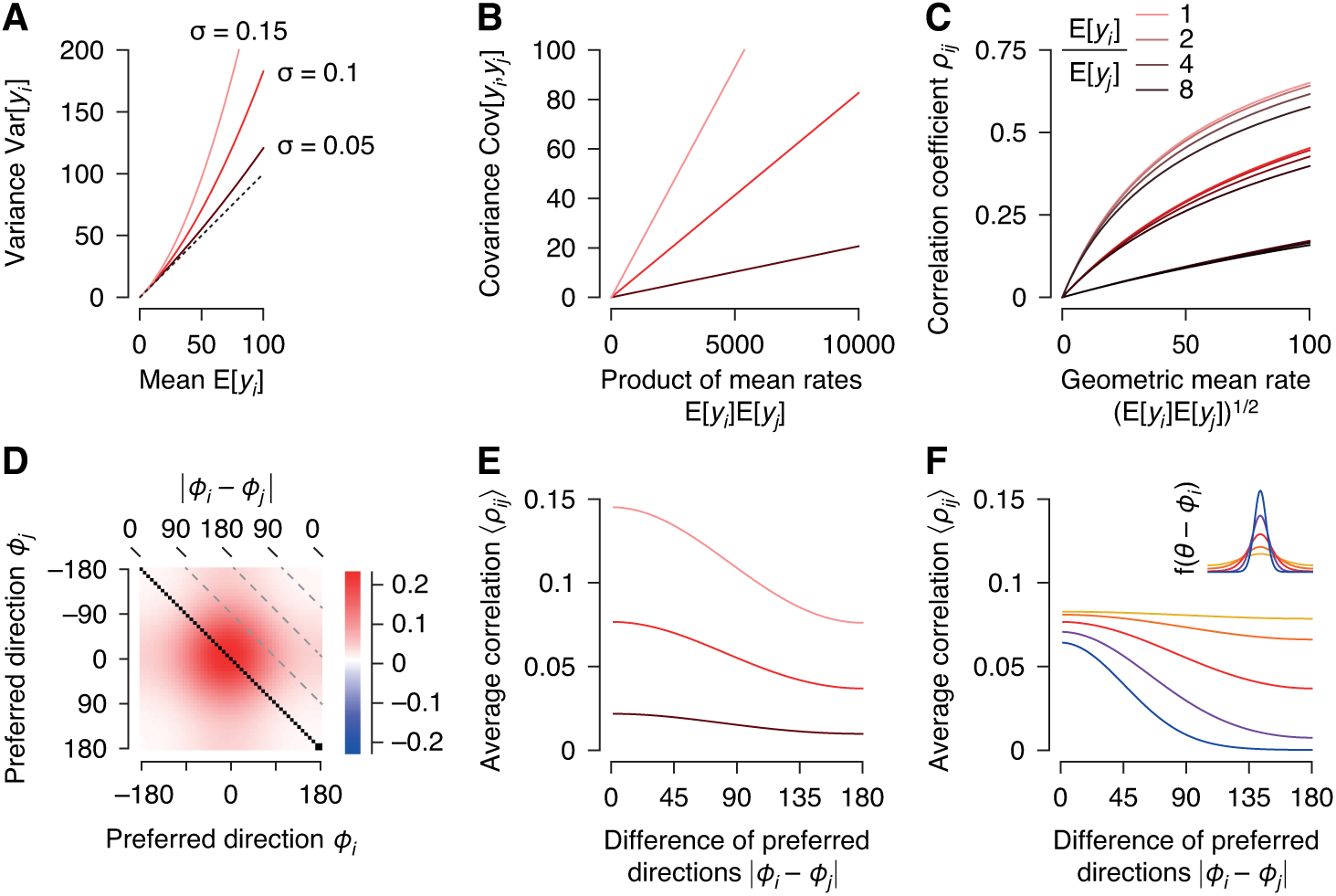
Effect of fluctuations in attentional state on spike count statistics. Solid lines: analytical solutions (Eqs. 31–35). Parameter values used here were *μ* = 0.1, *σ* ∈ {0.05, 0.1, 0.15} (dark to light red). **A.** Spike count variance as a function of mean spike count. Dashed line: identity (Poisson process). **B.** Covariance as a function of product of spike counts. **C.** Correlation coefficient as function of geometric mean firing rate. The three groups of lines correspond to different levels of *σ* as in the other panels. Darker colors within a group indicate increasing ratios *f*_*i*_(*θ*)*/f_j_*(*θ*). **D.** Matrix of correlation coefficients for *θ* = 0° and *σ* = 0.1. Tuning curves: *f*_*i*_(*θ*) = exp(*κ* cos(*θ* - *Φ*_*i*_) + *∈*), *κ* = 2, average firing rate across all *θ*: 10 spikes/s. **E.** Average correlation coefficient (over all directions of motion *θ*) as a function of difference of the preferred directions of the two neurons. Despite a common gain for all neurons, correlations decay with tuning difference. Parameters as in panel E. **F.** As in panel E, but for different tuning widths (*κ* ∈ {0.5, 1, 2, 4, 8}, shown in inset at the top). The decay of the correlations with the difference of the preferred directions is stronger for narrow tuning curves. Red line corresponds to panels D and E. Mean firing rate: 10 spikes/s for all tuning widths.

The first term is equal to the average spike count and results from the Poisson process assumption, while the second term is quadratic in the firing rate, which results from the multiplicative nature of the fluctuating gain *α* (Goris et al. 2014). Such an expanding mean-variance relation has been observed in many experimental studies (Britten et al. 1993; Dean 1981; Goris et al. 2014; Tolhurst et al. 1983). Note that if the attentional gain does not fluctuate, we recover the Poisson process.

Similar to the variances, we can compute the covariance between two neurons, which is given by the product of the firing rates and the variance of the attentional gain (Fig. 2B):

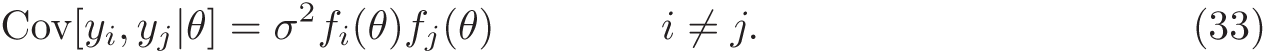

Recall that neurons are assumed to be conditionally independent given the attentional gain. Thus, any covariability arises exclusively from gain fluctuations. As a result, the covariance matrix (Fig. 2D) can be expressed as a diagonal matrix plus a rank-one matrix:

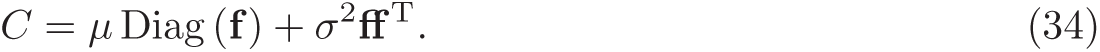

Note that the assumption of conditional independence could be relaxed without affecting any of the major results qualitatively: the diagonal matrix in the equation above would simply be replaced by the (non-diagonal) point process covariance matrix.

Experimental studies more typically quantify spike count correlations rather than covariances. We therefore also calculated the correlation coefficient *ρ*_*ij*_ of two neurons (Fig. 2C):

The spike count correlations induced by a fluctuating attentional gain increase with firing rates *f*_*i*_(*θ*). This effect, which has also been observed in numerous experimental studies (Cohen and Maunsell 2009; Ecker et al. 2014; Mitchell et al. 2009; Smith and Sommer 2013), arises because the independent (Poisson) variability is linear in the firing rate, whereas the covariance induced by gain fluctuations is quadratic and therefore dominates for large firing rates. Thus, correlations increase with the geometric mean firing rate, but there is no simple one-to-one mapping between the two quantities (it also depends on the ratio of the firing rates, Fig. 2C). The covariance, in contrast, is proportional to the product of the firing rates with a constant of proportionality of *σ*^2^ (Fig. 2B), suggesting that the latter might be more appropriate to consider when analyzing experimental data.

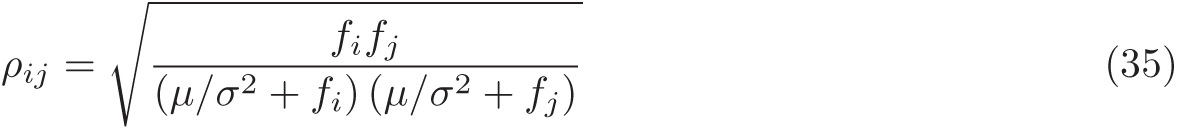

In addition, the correlation structure induced by gain fluctuations is non-trivial even if all neurons share the same gain (Fig. 2E, F; see also Ecker et al. (2014)). Due to the nonlinear shape of the tuning function and the nonlinear way the neurons’ tuning functions affect spike count correlations, the correlations decrease with increased difference in two neurons’ preferred directions (Fig. 2F). The slope of the decay depends mainly on the dynamic range of the tuning curve. If neurons have a high baseline firing rate compared to their peak firing rate, correlations decrease only marginally with preferred direction. In contrast, sharply tuned neurons with close to zero baseline firing rates exhibit strong limited-range structure.

This limited-range correlation structure has been observed in numerous experimental studies (Bair et al. 2001; Cohen and Maunsell 2009; Ecker et al. 2010; Smith and Kohn 2008; Zohary et al. 1994) and has been hypothesized to reflect shared input among similarly tuned neurons. However, our simple model shows that these seemingly structured correlations can arise from a very simple, non-specific mechanism: a common fluctuating gain that drives all neurons equally, irrespective of their tuning properties.

### 3.2 Fluctuations of feature attention

Feature attention is different from spatial attention in that the sign of the gain modulation depends on the similarity of the attended direction to the neuron’s preferred direction of motion (Fig. 3). Following the feature-similarity gain model (Treue and Martinez-Trujillo 1999), we model feature attention by

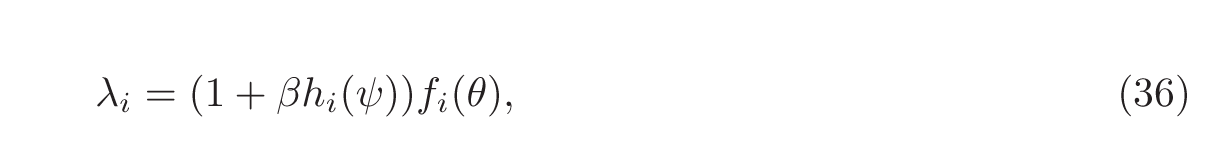

where *β* is the *feature gain* that controls how strongly the feature *Ψ* (in this case direction of motion) is attended on the given trial and *h*_*i*_(*Ψ*) is the *gain profile* (Fig. 3B) that determines the sign and relative strength of modulation for each neuron depending on the similarity of its preferred direction *Φ*_*i*_ to the attended direction *Ψ*. We assume that *h*_*i*_(*Ψ*) most strongly enhances neurons with preferred directions equal to the attended direction and suppresses those with opposite preferred directions (Fig. 3B).

**Figure 3:**
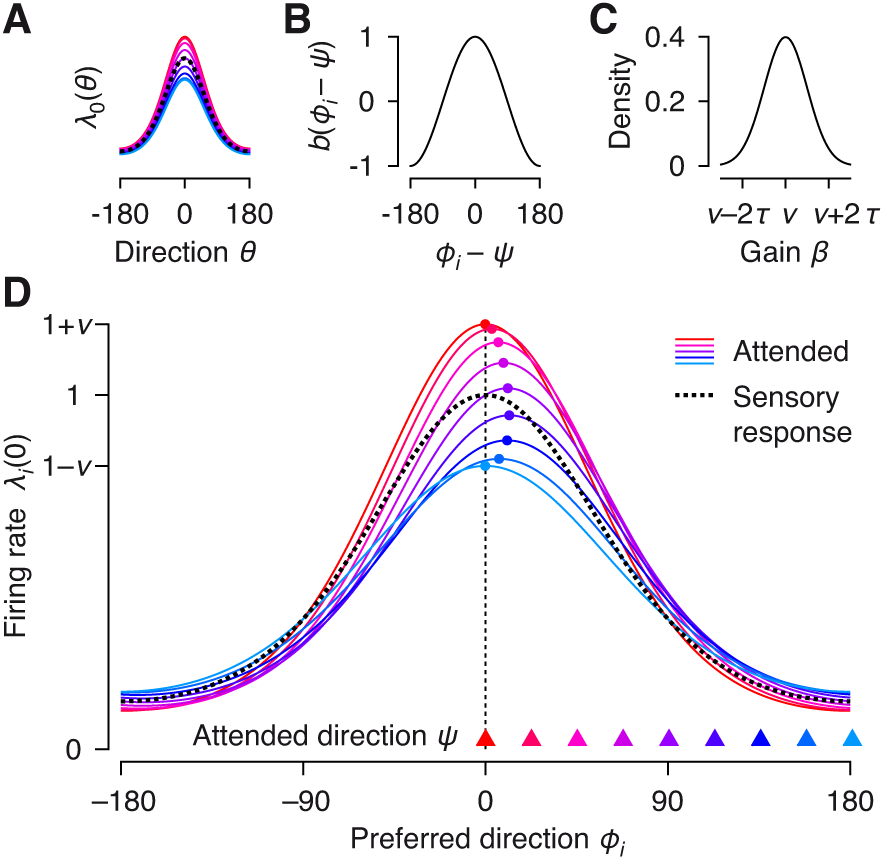
Model of feature attention. **A.** Tuning curve of a single neuron under sensory stimulation (black dotted) and with feature attention directed to different directions ranging from preferred (red) to null (blue). Note that the entire tuning curve of the neuron is gain-modulated and the modulation does not depend on the stimulus *θ*. **B.** The gain of a neuron depends on which direction of motion *Ψ* is attended relative to the neuron’s preferred direction *Φ*_*i*_. **C.** Distribution of gain (*β*) fluctuations. A Gaussian is shown for illustration purposes; the analysis holds for any distribution with E[*β*] = *ν* and Var[*β*] = *τ* ^2^. **D.** Population response of a homogeneous population of neurons under sensory stimulation (black dotted) and with attention directed to different directions of motion ranging from 0° (red) to 180° (blue). The stimulus is *θ* = 0. The curves show the average response of the neurons as a function of their preferred direction. Attending to a direction of motion biases the population response towards this attended stimulus. While each neuron’s tuning curve is gain-modulated as a whole (panel A), the population response is no longer equal to the individual neurons’ tuning curves, but instead sharpened/broadened and its peak is moved.

Because feature attention both increases and decreases different neurons’ gain depending on their preferred direction relative to the attended direction of motion, it biases the population response towards the attended direction (Fig. 3D). Thus, unlike in the case of spatial attention the shape of the population response is no longer identical to that of the individual neuron’s tuning curve. We start by assuming that the subject always attends the same direction (i. e. *Ψ* is constant) and consider the effect of fluctuations in the strength of attention, that is the gain *β*. We will come back to fluctuations in the attended direction below.

Similar to spatial attention, fluctuations in feature attention lead to overdispersion of the spike counts relative to a Poisson process (because rate variability is added).

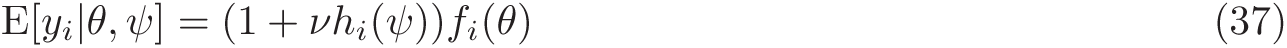

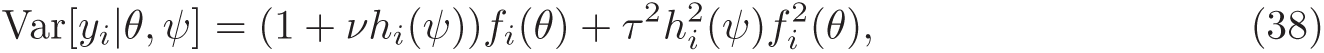

where *ν* = E[*β*] and *τ* ^2^ = Var[*β*] are the mean and the variance of the feature attention gain, respectively. The degree of overdispersion not only increases with the neuron’s firing rate, but also depends on the neuron’s preferred direction relative to the attended direction (Fig. 4A). Interestingly, spike counts are more overdispersed at the null direction than at the preferred direction (Fig. 4A: compare blue vs. black and green vs. yellow). The Fano factor (variance/mean) is given by

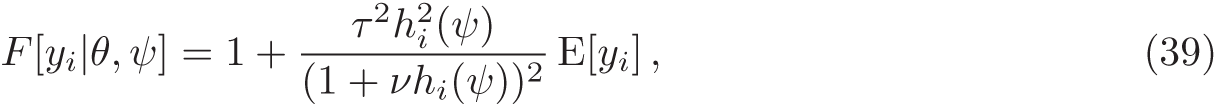

which is higher when *h*_*i*_ is negative than when it is positive. Neurons with preferred directions orthogonal to the attended direction are not overdispersed since *h*_*i*_ = 0.

**Figure 4:**
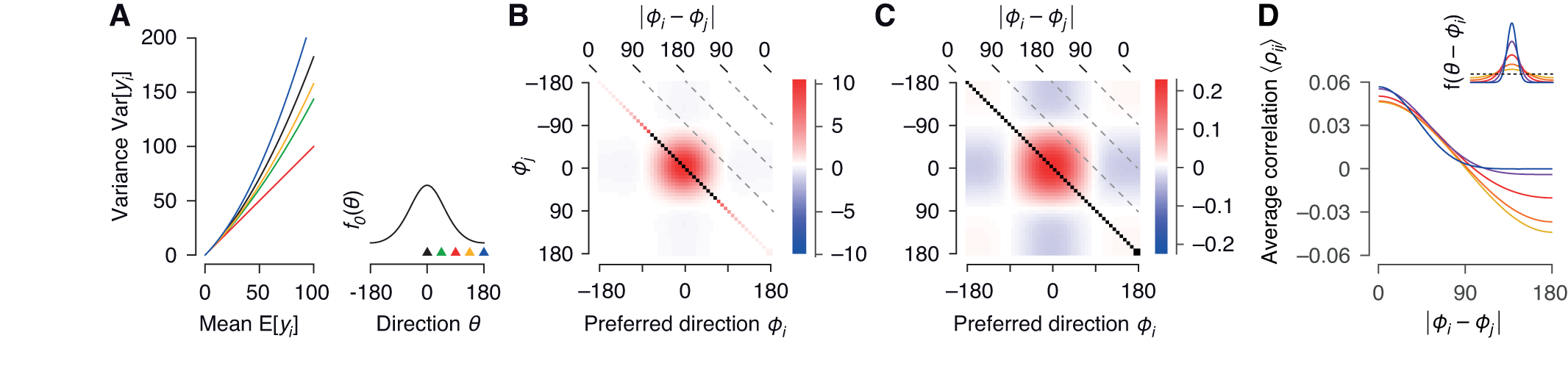
Effect of fluctuations in the feature attention gain on spike count statistics. Parameters here are: *Ψ* = 0, *ν* = 0.1, *τ* ^2^ = 0.01. **A.** Spike count variance as a function of mean spike count. Colors indicate different attended directions relative to the neurons’ preferred direction (*Φ*_*i*_ - *Ψ*; illustrated by colored triangles in inset on the bottom right). **B.** Covariance matrix for stimulus *θ* = 0. Neurons are ordered by preferred directions. Mean firing rate across the population: 20 spikes/s. **C.** As panel B, but the correlation coefficient matrix is shown. **D.** Dependence of spike count correlations on tuning similarity (difference of preferred directions). Fluctuations in feature attention induce limited range correlations irrespective of the shape of the tuning curve. The higher the baseline firing rate the stronger the negative correlations for neurons with opposite preferred directions. Inset: different tuning widths used.

As feature attention induces both increases as well as decreases in neuronal gain, the induced correlation structure is different from that induced by spatial attention. For the covariances, we obtain

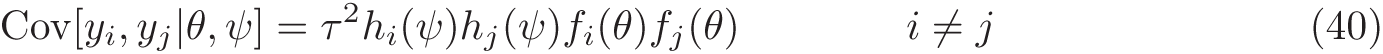

The sign of the covariance is determined by the product of *h*_*i*_ and *h*_*j*_, which depends on the attended direction and the preferred directions of the two neurons (Fig. 4B). For two neurons with identical preferred directions, the covariance is always positive while for two neurons with orthogonal preferred directions it is always negative. For any pair of neurons in between, it can be both positive and negative, depending on the stimulus (Fig. 4B). Again, the covariance matrix can be written as diagonal plus rank one:

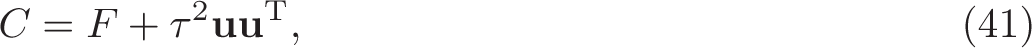

where *F*_*ii*_ = (1 + *νh_i_*(*Ψ*))*f*_*i*_(*θ*) and *u*_*i*_ = *h*_*i*_(*Ψ*)*f*_*i*_(*θ*).

As for spatial attention, averaging correlations over multiple stimulus conditions to represent the correlation structure as a function of the neurons’ tuning similarity misses much of the underlying structure (Fig. 4C): spike count correlations are positively correlated with tuning similarity (Fig. 4D), but the stimulus dependence (Fig. 4C) is again ignored. As before, the exact shape of the decay depends on the tuning width: for narrow tuning curves, neurons with opposite preferred directions are only weakly anti-correlated, whereas for broad tuning curves, those neurons are strongly anti-correlated (Fig. 4D, blue to red lines).

So far we have assumed that the attended direction of motion is constant and only the strength of attention fluctuates from trial to trial. Now we turn to the case where the attended direction fluctuates from trial to trial. We assume that, on average, the subject attends the correct direction, i. e. E[*Ψ*] = *θ*, but with some variance Var[*Ψ*] = *q*^2^. We further assume the gain *β* is constant. In this case, means and covariances of the observed spike counts are given by

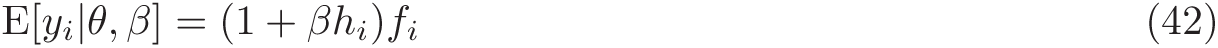

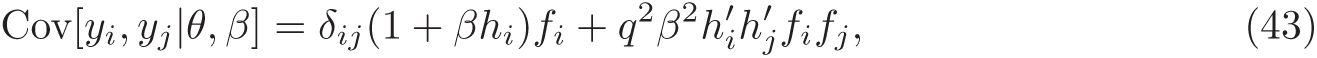

where 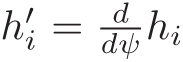 and we have abbreviated *h*_*i*_ ≡ *h*_*i*_(*θ*) and *f*_*i*_ ≡ *f*_*i*_(*θ*). As before, we can write the covariance matrix as diagonal plus rank one:

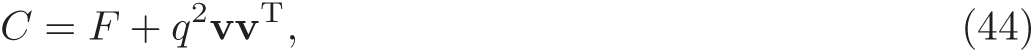

where *F*_*ii*_ = (1 + *βh_i_*)*f*_*i*_ and 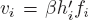. This pattern of correlations (Fig. 5) differs from those observed before for gain fluctuations in an important way: the sign of the correlation between two neurons depends only on whether their preferred directions are on the same side (both clockwise or counter-clockwise) of the stimulus direction or on different sides. As we will show more formally in the next section, this pattern of correlations is known as *differential correlations* (Moreno-Bote et al. 2014). Again, when plotted as a function of the difference of two neurons’ preferred directions, the correlations exhibit the typical limited-range structure (Fig. 5C), except for very narrow tuning curves, where the correlations are minimal around pairs with orthogonal preferred directions (Fig. 5C, blue lines). Also note that these correlations are substantially weaker than those induced by gain fluctuations (Figs. 2, 4), despite a relatively wide distribution of attended directions (SD: 10°).

**Figure 5:**
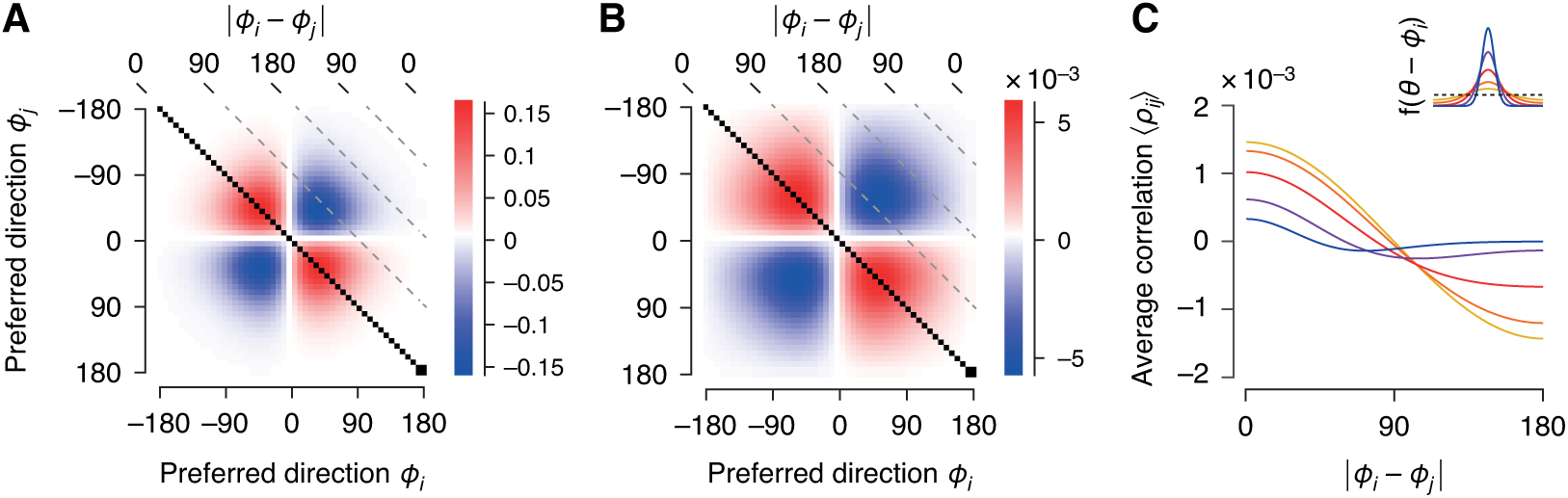
Effect of fluctuations in the attended direction on correlation structure. Parameters here are: E[*Ψ*] = 0, *q* = 10°, *β* = 0.1, *θ* = 0, mean firing rate across the population: 20 spikes/s. **A.** Covariance matrix. Neurons are ordered by preferred directions. **B.** As panel A, but the correlation coefficient matrix is shown. **C.** Dependence of spike count correlations on tuning similarity (difference of preferred directions). Fluctuations in the attended direction induce limited range correlations, whose shape depends on the width of the tuning curves. Inset: different tuning widths used.

### 3.3 Effect of attention-induced correlations on population coding

How interneuronal correlations affect the representational accuracy of neuronal populations has been a matter of immense interest (and debate) over the last years. Thus, we want to briefly consider how correlations induced by attentional fluctuations affect the coding accuracy of a population code.

Before doing so we need to make a choice: does the downstream readout have access to the state of attention or not? If it does, the picture is fairly simple: attentional fluctuations do not affect the readout accuracy, since the attentional state can be accounted for and there is no additional noise compared with a scenario without attentional fluctuations. The only downside is a potentially more complex readout. In contrast, if we assume that the readout does not have access to the attentional state, the situation becomes more interesting. In this case the attentional fluctuations act like additional (internally generated) noise, which could impair the readout. In the following we consider this latter scenario.

To quantify the accuracy of a population code, we use the Fisher information (Kay 1993) with respect to direction of motion. The Fisher information is useful because it quantifies the amount of information in a population of neurons without assuming a specific decoder. For a population of independent neurons, the Fisher information is linear in the number of neurons.

We start by considering spatial attention. Since the gain is the same for all neurons, gain fluctuations should not affect the coding accuracy of the population with respect to the direction of the stimulus, which is encoded in the differential activation pattern of the neurons. This is indeed the case. The Fisher information of a population of Poisson neurons whose firing rates are modulated by a common gain with mean *μ* is given by

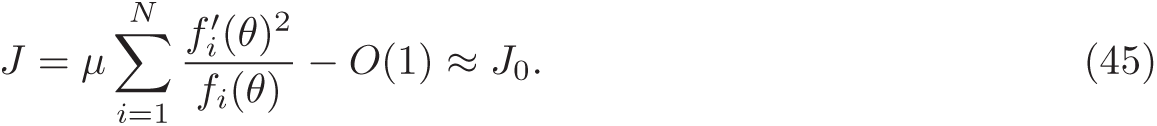

Thus, unobserved gain fluctuations reduce the information in the population only by a constant term (for derivation see Appendix). For reasonably large populations (e.g. *N >* 100) this term can be neglected and the information is approximately equal to that of an independent population (*J*_0_). This result can be understood intuitively by considering the structure of the covariance matrix (Eq. 33): the dominant eigenvector points in the direction of the tuning function f, which is orthogonal to changes in the stimulus, f ^’^. Therefore, gain flucutations do not impair the readout of the direction of motion.

The same result holds for fluctuations in the feature attention gain, so long as the attended direction matches the one shown and does not fluctuate from trial to trial. A fluctuating gain sharpens and broadens the population hill from trial to trial, but leaves its peak unchanged. Again, the dominant eigenvector (*u*_*i*_ = *h*_*i*_*f*_*i*_, Eq. 40) points in a direction that is orthogonal to changes in the stimulus (details see Appendix).

The situation changes if the focus of attention (i. e. the attended direction) fluctuates from trial to trial or the attended direction does not match the one shown: since feature attention biases the population response towards the attended direction, such attentional fluctuations have the same effect as noise on the input [*differential correlations*, (Moreno-Bote et al. 2014)]. To illustrate this finding, we switch to a slightly modified and more specific response model than above. Assuming *f*_*i*_(*θ*) = exp(*κ* cos(*θ* - *Φ*_*i*_)) and *h*_*i*_ = cos(*Ψ* - *Φ*_*i*_), and noting that (1 + *βh_i_*) ≈ exp(*βh_i_*), we can write the log-firing rate as

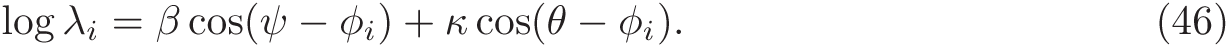

We can combine the two cosine terms and obtain:

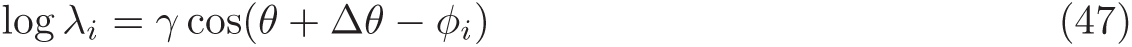

Where

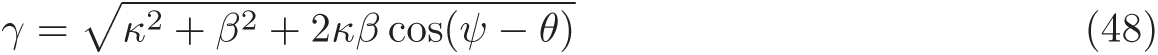

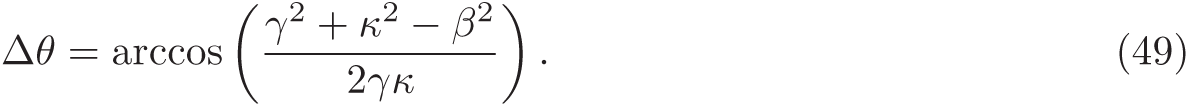

Thus, feature attention biases the population response away from the stimulus direction *θ* towards the attended direction *Ψ*. The magnitude of the bias Δ*θ* depends on both the strength of feature attention *β* and the attended direction *Ψ*. Consequently, if *Ψ* ≠ *θ* fluctuations in either the attended feature or the degree of feature attention have the same effect on the population response as variance of the stimulus direction that is shown, i. e. they induce differential correlations. This result can also be understood by considering the structure of the covariance matrix (Eq. 44): the dominant eigenvector 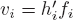 points in the same direction as changes in the stimulus, f’. We can therefore approximate the Fisher information by (see Moreno-Bote et al. 2014)

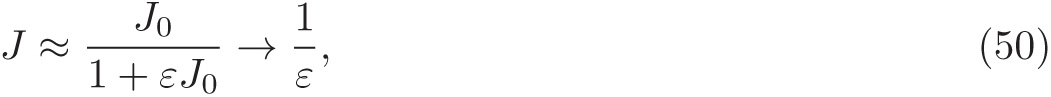

where *J*_0_ is again the information in an independent population and *ε* = Var[Δ*θ*] depends on both the distribution of attended directions and the variance of the gain. In this case, the information in the population saturates at a finite value 1*/ε* that depends only on the distribution of the attention signal and can be substantially lower than the limit imposed by the information in feedforward signal (see also Discussion). When the subject attends the correct direction on average (i. e. E[*Ψ*] = *θ*) and the variance of the attended direction (Var[*Ψ*]) is small, we find

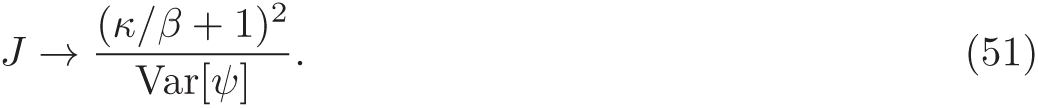

Thus, the saturation level depends on the strength (*β*) of attention relative to the tuning width (*κ*) and the variance in the attended direction.

### 3.4 Identifying attentional fluctuations in experimental data

We saw above that fluctuations in attentional state can introduce interesting patterns of correlations in neural activity, all of which are roughly consistent with the published literature on attention. However, as long as one considers only single neurons and pairwise statistics, any result can be consistent with many hypotheses. For instance, attentional fluctuations induce correlations that depend on firing rates (Fig. 2C), but the same result is also predicted by the thresholding non-linearity of neurons (Rocha et al. 2007) and therefore need not result from attentional fluctuations. Similarly, all types of attentional fluctuations considered above lead to correlations that decrease with the difference of two neurons’ preferred directions (*limited range correlations*, Figs. 2E, 4D, 5C), but this correlation structure can also arise from shared sensory noise (Shadlen and Newsome 1998).

So how would one go about identifying attentional fluctuations in experimental data? Clearly, one has to consider the response patterns of simultaneously recorded populations of neurons rather than just pairwise correlations. In the following, we discuss some predictions our model makes for the structure of the neural population response.

A first approach suggested by our analyses above: we showed that in all cases we analyzed the covariance matrix induced by attentional fluctuations is diagonal plus rank one. Thus, attentional fluctuations are restricted to a low-dimensional subspace that could be identified from simultaneously recorded neurons by *Factor Analysis*. However, the disadvantage of this approach is that the low-dimensional subspace depends on the stimulus in a non-trivial way (Fig. 6A for *θ* = 0° [left] and *θ* = 60° [right]; see also Eqs. 33, 40, 44). This stimulus dependence precludes pooling of data over multiple stimulus conditions. Moreover, if the attended direction does not match the stimulus direction, the major axes of variability do not peak at either direction, but somewhere in between (Fig. 6A, blue lines in the right panel, where *Ψ* = 0° and *θ* = 60°). Thus, it is non-trivial to recover the quantities of interest for the experimenter – the attended feature (direction) and the degree of attention allocated (the gain).

**Figure 6:**
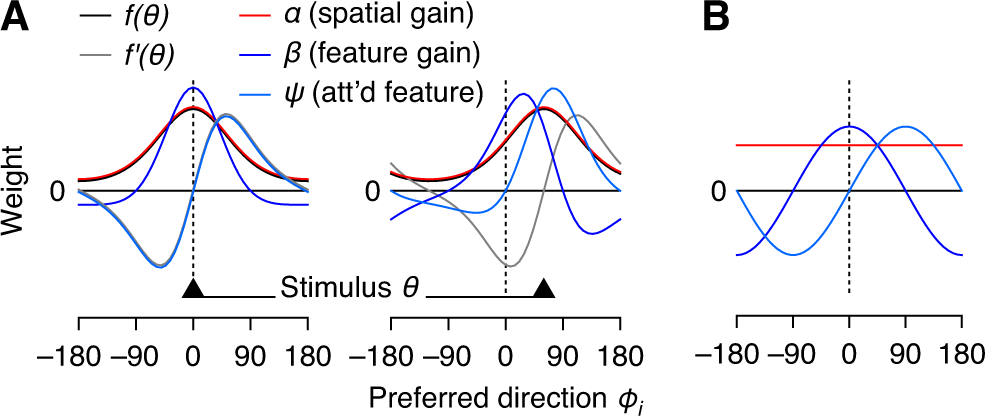
Identifying attentional fluctuations from variability in neuronal population activity. **A.** The subspace identified by Factor Analysis depends on the stimulus direction. Black triangles: stimulus direction (left: *θ* = 0°, right: *θ* = 60°). Solid lines: basis functions corresponding to fluctuations in spatial attention gain (red), feature attention gain (dark blue), and attended direction (light blue); population tuning curve (black) and its derivative (gray). Horizontal dashed line: (average) attended direction. **B.** Principal components identified by Exponential Family PCA are independent of the stimulus since the log-link turns a multiplicative modulation into an additive offset. Colors as in panel A.

A model that could directly extract attentional gains (spatial and feature gain) and the attended feature would be desirable. Fortunately, all three can be inferred from population activity in a straightforward manner using methods such as *Exponential Family Principal Component Analysis (E-PCA)* (Collins et al. 2001; Mohamed et al. 2009) or *Poisson Linear Dynamical Systems (PLDS)* (Buesing et al. 2012; Macke et al. 2011). Similar to above (Eq. 46), we assume *f*_*i*_(*θ*) = exp(*κ*_*i*_ cos(*θ*- *Φ*_*i*_) + *ε*_*i*_) and *h*_*i*_ = cos(*Ψ* - *Φ*_*i*_) and write the log-firing rate as

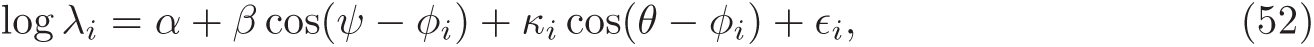

which can be rewritten as a linear function of the attentional state and the stimulus:

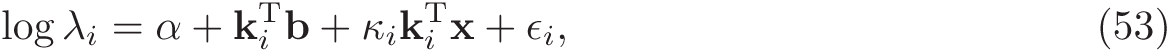

where *α* and b = *β*.[cos *Ψ,* sin *Ψ*]^T^ represent the state of spatial and feature attention, respectively, x = [cos *θ,* sin *θ*]^T^ is the stimulus, *k*_*i*_ = [cos *Φ_i_,* sin *Φ*_*i*_]^T^ is the neuron’s preferred direction, *κ*_*i*_ the (inverse) tuning width, and *ε*_*i*_ controls the mean firing rate. This model is a *Generalized Linear Model (GLM)* with Poisson observations and log(*x*) as the link function. Thus, E-PCA or PLDS will recover the subspace corresponding to fluctuations in attentional state {*α,* b}. This subspace is spanned by *u*_*i*_ = [1, cos *Φ_i_,* sin *Φ*_*i*_] and independent of the stimulus (see Fig. 6B). The attentional gains are *α* and *β* = ||b||, while the attended direction is *Ψ* = ∠b.

### 3.5 A new view on the reduction of shared variability under attention

There is ample experimental evidence that attention fluctuates from trial to trial (Cohen and Maunsell 2010; Cohen and Maunsell 2011), and we showed in the previous sections that such fluctuations induce patterns of (correlated) variability that are highly consistent with the reported data on attention (Cohen and Maunsell 2009; Herrero et al. 2013; Mitchell et al. 2009). Interestingly, in our model, both the magnitude of overdispersion in single neurons’ spike counts and the average level of correlations are determined by the variance of the attentional gain (*σ*^2^ = Var[*g*]), but not by its average modulation (*μ* = E[*g*]). This observation suggests that the average attentional modulation (*μ*) between an attended and an unattended condition (which can be reliably measured based on average responses) does not predict the level of correlations in either condition, since the latter is controlled by an independent variable (*σ*^2^). Indeed, this is one of the central experimental findings: directing spatial attention to a certain location *increases* the average responses of neurons whose receptive fields represent this location, but reduces independent and shared variability among those neurons (Cohen and Maunsell 2009; Herrero et al. 2013; Mitchell et al. 2009). Thus, if our model is correct, then the data suggest that attention not only increases response gain, but also reduces the trial-to-trial variability of the gain.

This view of attention has important implications for the role of interneuronal correlations under attention. Recent studies (Cohen and Maunsell 2009; Mitchell et al. 2009) have argued that spatial attention improves behavioral performance primarily by reducing correlations. However, as we showed above, fluctuations of spatial attention do not affect the representational accuracy of the neuronal population. Therefore, under our model the experimentally observed reduction in correlations is irrelevant when reading out a neuronal population. The only difference that matters is the increase in gain.

## 4 Discussion

We find that a simple model of neuronal responses can account for a range of empirically observed phenomena relating attention, neuronal variability and coding properties of neuronal populations. Our model unites two central findings in the literature on attention, that attention acts as a multiplicative gain factor on neuronal responses (Maunsell and Treue 2006) and that attention fluctuates from trial-to-trial (Cohen and Maunsell 2010). The importance of the combined effects of these observations has not previously been fully appreciated. We show that such a model is sufficient to account for super-Poisson variability (see also Ecker and Tolias 2014; Goris et al. 2014) as well as a variety of pairwise correlation structures, most notably the limited-range structure and differential correlations (Abbott and Dayan 1999; Ecker et al. 2010; Moreno-Bote et al. 2014; Smith and Kohn 2008).

Our results argue that it is likely that a large fraction of variability in the neuronal response can be attributed to fluctuations in behaviorally relevant, internally-generated signals, such as attention, rather than shared noise (Ecker and Tolias 2014; Ecker et al. 2010, 2014; Goris et al. 2014; Nienborg and Cumming 2009). This view suggests the hypothesis that correlations that arise from such fluctuating signals generally should not impair coding of sensory information. We find that this assertion is true for the case of fluctuations in the magnitude of the gain. The Fisher information of our model population of neurons is not limited by fluctuations in the strength of attentional gain (i.e., is independent of the variance of the gain term), despite those fluctuations generating a limited-range correlation structure typically thought to impair coding.

However, theoretical work has shown that the effect of different patterns of correlations on the coding of sensory information is nuanced and can depend greatly on specific assumptions that are made regarding a variety of neuronal properties, such as the shapes of tuning curves in the population, subtle details of the assumed correlation structure, or different readouts (Abbott and Dayan 1999; Ecker et al. 2011; Josić et al. 2009; Shamir and Sompolinsky 2006; Sompolinsky et al. 2001; Wilke and Eurich 2002). The recent work of Moreno-Bote et al. (2014) has helped to clarify the problem of when and what types of correlation structures are detrimental to coding with their description of differential correlations, a specific pattern of correlation proportional to the product of the derivative of the tuning curves that leads to information saturation. Our model generates this pattern of correlated variability when the fluctuations in attention occur around a specific feature rather than a specific gain value. Thus, it is noteworthy that a model only slightly more complicated than typical Poisson spiking models can generate the diversity of correlation structures noted in the experimental and theoretical literature as being important for population coding.

In addition to offering a parsimonious account of neuronal variability and co-variability, our model has implications for how we should interpret the effect of attention as it relates to improvements in perceptual performance. Chiefly, if the reduction of correlations observed under attention is indeed due to a reduction of gain fluctuations – as our model would suggest – the reduction of correlations is irrelevant with respect to the coding accuracy of the population and cannot be the mechanism improving behavioral performance as suggested by recent experimental studies (Cohen and Maunsell 2009; Herrero et al. 2013; Mitchell et al. 2009).

Our model leads to a second interesting observation: It is likely that not only the attentional gain fluctuates from trial to trial, but also the attended feature itself. Such fluctuations introduce differential correlations, which indeed impair the readout (unless it has exact access to the attended feature). Thus, the attentional mechanism itself places a limit on how accurately a stimulus can be represented by a sensory population, and this limit can at least in principle be substantially lower than the amount of sensory information entering the brain through the eye. This insight may trigger the question: why, then, should there be an attentional mechanism in the first place? There are a number of possible answers to this question.

First, we can think of attention as a prior. Using prior information to bias an estimate towards more likely solutions will on average improve the estimate. In situations where the stimulus is noisy and decisions have to be made fast, such a bias is most beneficial and outweighs the small extra noise added due to variability in the prior. Conversely, in situations where there is lots of sensory evidence, the full information content present in the eye is rarely necessary in real-world situations, and, therefore, the noise added due to attentional fluctuations does not matter either.

Second, it should be noted that for change-detection paradigms that are typically employed in attention experiments, the estimation framework that asks how well a stimulus value can be reconstructed (e. g. Fisher information) is not quite appropriate. In such tasks the subject never judges the *absolute* direction (or any other feature) of the stimulus, but instead has to detect a small change, that is the difference between two subsequent stimuli. In this case any errors introduced due to fluctuations in the attended direction cancel out, since they affect both stimuli roughly equally, at least so long as attentional fluctuations occur at a timescale that is slow enough, such that the attentional state is approximately the same for both the pre-and post-change stimulus.

## References

Abbott, L. F. and P. Dayan (1999). “The Effect of Correlated Variability on the Accuracy of a Population Code”. Neural Computation 11.1, pp. 91–101.

Bair, W., E. Zohary, and W. T. Newsome (2001). “Correlated Firing in Macaque Visual Area MT: Time Scales and Relationship to Behavior”. The Journal of Neuroscience 21.5, pp. 1676–1697.

Britten, K. H. et al. (1993). “Responses of neurons in macaque MT to stochastic motion signals”. Visual Neuroscience 10.06, pp. 1157–1169.

Buesing, L., J. H. Macke, and M. Sahani (2012). “Learning stable, regularised latent models of neural population dynamics”. Network: Computation in Neural Systems 23.1-2, pp. 24–47.

Cohen, M. R. and J. H. R. Maunsell (2009). “Attention improves performance primarily by reducing interneuronal correlations”. Nature Neuroscience 12.12, pp. 1594–1600.

Cohen, M. R. and J. H. R. Maunsell (2010). “A Neuronal Population Measure of Attention Predicts Behavioral Performance on Individual Trials”. The Journal of Neuroscience 30.45, pp. 15241– 15253.

Cohen, M. R. and J. H. Maunsell (2011). “Using Neuronal Populations to Study the Mechanisms Underlying Spatial and Feature Attention”. Neuron 70.6, pp. 1192–1204.

Collins, M., S. Dasgupta, and R. E. Schapire (2001). “A generalization of principal components analysis to the exponential family”. In: Advances in neural information processing systems, pp. 617–624.

Dean, A. F. (1981). “The variability of discharge of simple cells in the cat striate cortex”. en. Experimental Brain Research 44.4, pp. 437–440.

Ecker, A. S. and A. S. Tolias (2014). “Is there signal in the noise?” en. Nature Neuroscience 17.6, pp. 750–751.

Ecker, A. S. et al. (2010). “Decorrelated Neuronal Firing in Cortical Microcircuits”. Science 327.5965, pp. 584–587.

Ecker, A. S. et al. (2011). “The Effect of Noise Correlations in Populations of Diversely Tuned Neurons”. The Journal of Neuroscience 31.40, pp. 14272–14283.

Ecker, A. S. et al. (2012). “The correlation structure induced by fluctuations in attention”. In: Cosyne Abstracts, pp. III–46.

Ecker, A. S. et al. (2014). “State dependence of noise correlations in macaque primary visual cortex”. Neuron 82.1, pp. 235–248.

Goris, R. L. T., J. A. Movshon, and E. P. Simoncelli (2014). “Partitioning neuronal variability”. en. Nature Neuroscience 17.6, pp. 858–865.

Herrero, J. et al. (2013). “Attention-Induced Variance and Noise Correlation Reduction in Macaque V1 Is Mediated by NMDA Receptors”. Neuron 78.4, pp. 729–739.

Josić, K. et al. (2009). “Stimulus-Dependent Correlations and Population Codes”. Neural Computation 21.10, pp. 2774–2804.

Kay, S. M. (1993). Fundamentals of Statistical Signal Processing, Volume I: Estimation Theory. 1st ed. Prentice Hall.

Macke, J. H. et al. (2011). “Empirical models of spiking in neural populations”. Advances in neural information processing systems 24, p. 13501358.

Maunsell, J. H. and S. Treue (2006). “Feature-based attention in visual cortex”. Trends in Neurosciences 29.6, pp. 317–322.

Mitchell, J. F., K. A. Sundberg, and J. H. Reynolds (2009). “Spatial Attention Decorrelates Intrinsic Activity Fluctuations in Macaque Area V4”. Neuron 63.6, pp. 879–888.

Mohamed, S., Z. Ghahramani, and K. A. Heller (2009). “Bayesian exponential family PCA”. In: Advances in Neural Information Processing Systems, pp. 1089–1096.

Moreno-Bote, R. et al. (2014). “Information-limiting correlations”. en. Nature Neuroscience advance online publication.

Nienborg, H. and B. G. Cumming (2009). “Decision-related activity in sensory neurons reflects more than a neuron’s causal effect”. Nature 459.7243, pp. 89–92.

Rabinowitz, N. et al. (2015). “Modulators of V4 population activity under attention”. In: Cosyne Abstracts.

Reynolds, J. H. and L. Chelazzi (2004). “Attentional Modulation of Visual Processing”. Annual Review of Neuroscience 27.1, pp. 611–647.

Rocha, J. de la et al. (2007). “Correlation between neural spike trains increases with firing rate”. Nature 448.7155, pp. 802–806.

Shadlen, M. N. and W. T. Newsome (1998). “The Variable Discharge of Cortical Neurons: Implications for Connectivity, Computation, and Information Coding”. The Journal of Neuroscience 18.10, pp. 3870 –3896.

Shamir, M. and H. Sompolinsky (2006). “Implications of Neuronal Diversity on Population Coding”. Neural Computation 18.8, pp. 1951–1986.

Smith, M. A. and A. Kohn (2008). “Spatial and Temporal Scales of Neuronal Correlation in Primary Visual Cortex”. J. Neurosci. 28.48, pp. 12591–12603.

Smith, M. A. and M. A. Sommer (2013). “Spatial and Temporal Scales of Neuronal Correlation in Visual Area V4”. en. The Journal of Neuroscience 33.12, pp. 5422–5432.

Sompolinsky, H. et al. (2001). “Population coding in neuronal systems with correlated noise”. Physical Review E 64.5, p. 051904.

Tolhurst, D. J., J. A. Movshon, and A. F. Dean (1983). “The statistical reliability of signals in single neurons in cat and monkey visual cortex”. Vision research 23.8, pp. 775–785.

Treue, S. and J. C. Martinez-Trujillo (1999). “Feature-based attention influences motion processing gain in macaque visual cortex”. Nature 399.6736, pp. 575–579.

Wilke, S. D. and C. W. Eurich (2002). “On the functional role of noise correlations in the nervous system”. Neurocomputing 44-46, pp. 1023–1028.

Zohary, E., M. N. Shadlen, and W. T. Newsome (1994). “Correlated neuronal discharge rate and its implications for psychophysical performance”. Nature 370.6485, pp. 140–143.

